# Pericardial & Mediastinal Fat-Associated Lymphoid Clusters are rapidly activated in an alkane induced model of Systemic Lupus Erythematosus

**DOI:** 10.1101/2023.07.28.549766

**Authors:** Karolina Bentkowska, Alex Hardgrave, Nadia Iqbal, Laura Gresty, Bethany Marsden, Sheila Macharia, Lucy Jackson-Jones

## Abstract

Systemic Lupus Erythematosus (SLE) is an autoimmune disease predominated by auto-antibodies that recognise cellular components. Pleural involvement is the most common SLE-related lung disease. Natural antibodies are rapidly secreted by innate-like B cells following perturbation of homeostasis and are important in the early stages of immune activation. The serous cavities are home to large numbers of innate-like B cells present both within serous fluid and resident within fat-associated lymphoid clusters (FALCs). FALCs are important hubs for B-cell activation and local antibody secretion within the body cavities. Patients with SLE can develop anti-phospholipid antibodies and in rare situations develop alveolar haemorrhage. Utilising delivery of the hydrocarbon oil pristane in C57BL/6 mice as a model of SLE we identify a rapid expansion of pleural cavity B cells as early as day 3 after intra-peritoneal pristane delivery. Following pristane delivery, pericardial B1 B cells are proliferative, express the plasma-cell surface marker CD138 and secrete both innate and class switched antibodies highlighting that this cavity niche may play an unrecognised role in the initiation of lupus pleuritis.

**Graphical abstract:** 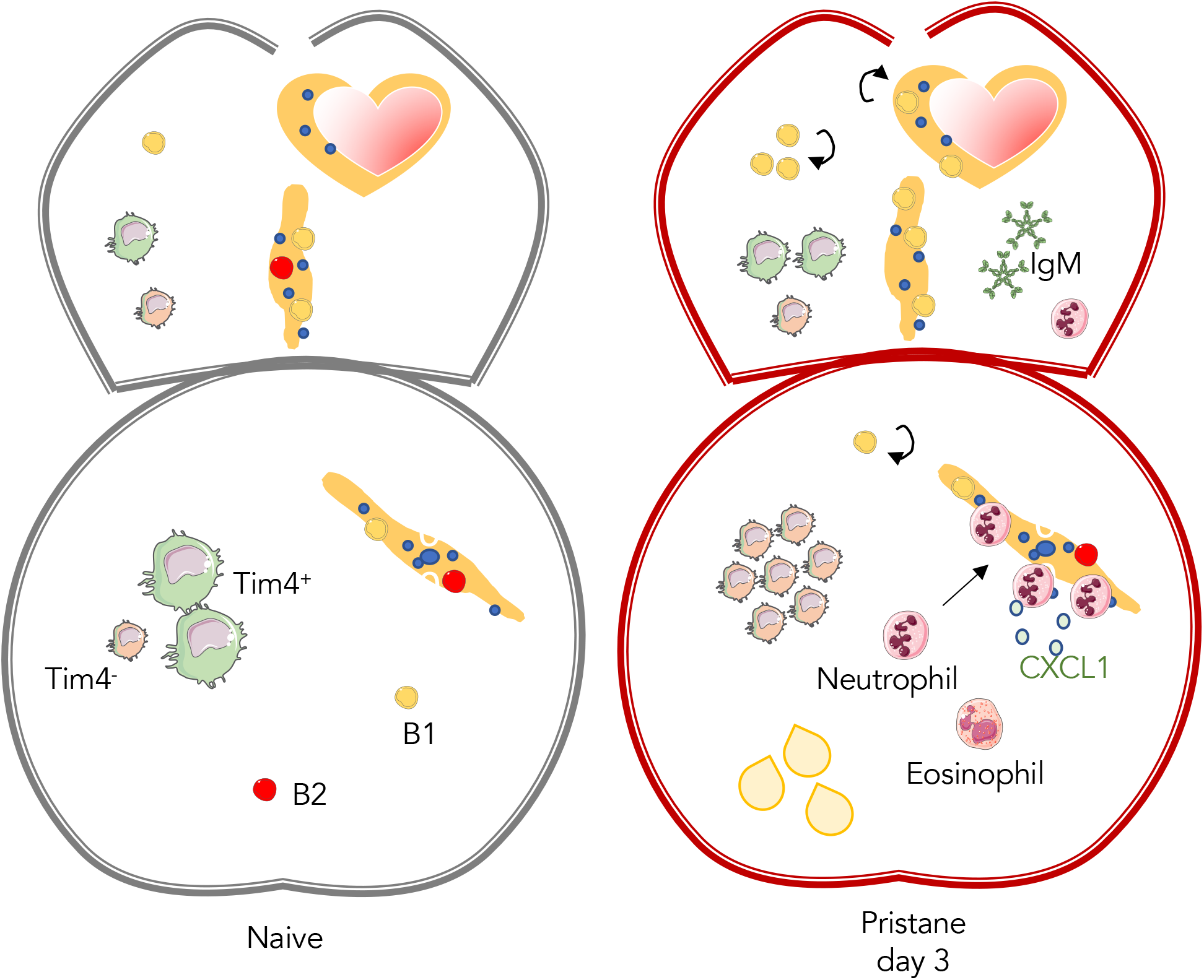

## Introduction

Systemic lupus erythematosus (SLE) is an auto-immune disease characterized by the production of systemic anti-nuclear antibodies, development of arthritis, immune complex-mediated glomerulonephritis as well as pericardial and pleural inflammation. The mechanism of pleural involvement in SLE is under-investigated however 45-60% of SLE patients experience pleuritic pain and pleural involvement is one of the most common SLE-lung related diseases(Pego-Reigosa, Medeiros, and Isenberg 2009). Overproduction of natural antibodies against self-antigens is a characteristic feature of autoimmune diseases such as systemic lupus erythematosus (Reyneveld et al., 2020). Furthermore, 30-40% of SLE patients are positive for the presence of anti-phospholipid antibodies (Unlu, Zuily, and Erkan 2016). Previous studies have also confirmed the presence of anti-nuclear antibodies within pleural fluid of patients with SLE and found these to be at a higher titer within pleural fluid than the serum suggesting a local accumulation or production (Good et al. 1983). The serous cavities, including the pleural cavity are predominated by B1 B cells (Finlay and Allen 2020) and are key sites at which natural antibodies are detected following their local secretion from fat-associated lymphoid clusters (FALCs) (Jones et al. 2015; Jackson-Jones et al. 2016). Omental FALCs (previously known as milky spots) are also important sites providing protection to the serous cavities in the context of inflammation (Benezech et al. 2015; Jackson-Jones 2020), infection (Zhang et al. 2019; Christian et al. 2022) and cancer (Etzerodt et al. 2020). The alkane tetramethylpentadecane (TMPD, commonly known as pristane) is a hydro-carbon oil which has been shown to model SLE following injection *in vivo* in animal models (Reeves et al. 2009).

Studies utilising pristane mainly focus on the detection of natural antibodies in the serum typically weeks after pristane delivery when the response becomes systemic (Pawar et al. 2014; Han et al. 2015; Zhuang et al. 2016; Liu et al. 2020; Yan et al. 2020). In different strains of mice, pristane induces distinct responses, for example BALB/c mice go on to develop arthritis whereas C57BL/6 mice instead develop pulmonary complications including diffuse alveolar haemorrhage (Reeves et al. 2009). The baseline number and activation of fat-associated lymphoid clusters within the pericardium and mediastinum of mice (Jackson-Jones et al. 2016) also differs by strain, with those on the C57BL/6 background being more strikingly activated during allergic inflammation than BALB/c. A role for IgM in the development of thoracic complications of SLE has been shown previously; B cell deficient μMT^-/-^ mice that are resistant to development of diffuse alveolar haemorrhage become susceptible following infusion of IgM (Zhuang et al. 2017). As FALCs within the pericardium and mediastinum are key sites for local IgM secretion within the pleural cavity (Jackson-Jones et al. 2016) and expansion of mediastinal FALCs has been shown in an MRL/MpJ-*lpr* autoimmune mouse model of SLE (Elewa, Ichii, and Kon 2016) we aimed to determine whether these structures are activated during pristane induced pleuritis.

Studies of pristane exposure commonly assess systemic auto-antibody responses in the weeks to months following exposure and as such, the early immune response within the pleural cavity remain relatively elusive. To better understand the pleural cavity response to pristane we compared the early events occurring at this site with those in the peritoneal cavity which is the site of pristane delivery. We found higher numbers of B-cells within the pleural cavity by day 3 following i.p. delivery of pristane with increased proliferation of B1 B cells within both pleural fluid and FALCs. In contrast to the omentum, which did not show increased antibody secretion, there was a significant increase in total IgM, IgG2a, & IgG1 antibody released by the pericardial and mediastinal FALCs at day 3 following exposure to pristane. Furthermore, we detected an increase in natural IgM antibodies that recognize phospho-lipids within lavage fluid from the pleural cavity following pristane delivery.

## Methods

### Animals

Experiments performed at Lancaster University were conducted in accordance with the Animals (Scientific Procedures) Act, UK 1986 under license granted by the Home Office (UK) following prior approval by local animal welfare and ethical review body. Experiments were performed using female C57BL/6J mice aged 8-12 weeks, animals were purchased from Charles River or were the wildtype offspring of genetically modified animals that were bred and maintained under specific pathogen–free conditions at Lancaster University.

### In vivo procedures

Mice were injected i.p. with 300μl of the hydrocarbon oil pristane (Sigma) or left naïve. Pleural & peritoneal exudate cells (PLEC & PEC) were isolated by flushing murine cavities with a minimum of 2ml of RPMI 1640 (Sigma). Cell pellets were isolated by centrifugation and the first 2 × 1ml of lavage fluid was stored at -20 to -80°C for further analysis. Omentum, pericardial and mediastinal adipose tissues were isolated following lavage, weighed and in select experiments omentum and mediastinal adipose were cultured ex vivo in 500μl of complete RPMI 1640 (sigma) containing 10% Fetal calf serum (Sigma), 1% v/v Penicillin Streptomycin (Gibco) for 2h at 37°C, 5% CO_2_ and the culture supernatant stored as above.

### Flow cytometry

Murine cells were stained to distinguish live versus dead using Zombie aqua (Biolegend), blocked with anti-murine CD16/32 (clone 2.4G2, Biolegend) & 10% rat & mouse serum (Sigma) and stained for cell surface markers (See Supplementary Table 1 for list of antibodies used), following staining cells were washed using FACs buffer (2% BSA, 2mM EDTA, PBS). In select experiment cells were fixed and permeabilised using eBioscience-FoxP3 fixation and permeabilization buffer and incubated with anti-Ki67 anti-bodies for a minimum of 30 minutes prior to washing with acquisition. All samples were acquired using a Beckman Coulter Cytoflex and analyzed with FlowJo software (Tree Star).

### Wholemount immunofluorescence staining

Murine omentum, pericardial and mediastinal adipose tissues were fixed for one hour on ice in 10% NBF (Fisher or Sigma) then permeabilized in PBS 1% Triton-X 100 (Sigma) for 20 minutes at room temperature prior to staining with primary antibodies for 1 hour in PBS 0.5% BSA 0.5% Triton (Sigma). Antibodies used are listed in Supplementary Table 1.

### Microscopy

After mounting with Fluoromount G (Invitrogen), confocal images were acquired using a Leica Stellaris 5 laser scanning confocal microscope or a Zeiss LSM880, analysis was performed using FIJI (Image, J). To calculate the area of FALCs stained with antibodies of interest, the perimeter of the FALC was delimited manually and a fixed threshold for IgM, F4/80 and Ki67 fluorescence was set.

### Detection of cytokines, chemokines & antibodies

A mix and match Murine anti-viral Legendplex array (Biolegend) was used to detect CXCL1 and a 6-plex murine isotyping panel was used to detect total IgM, IgG2a, IgG1 and IgA within tissue culture supernatants following the manufacturer’s instructions with the modification that half of the suggested beads and reagents were used per assay; following prior optimisation. For the detection of IgM recognising phospholipid species, 5μg/ml of oxLDL (Invitrogen), POVPC (Avanti lipids), OxPAPC (Avanti lipids) were coated overnight on 96-well high-binding ELISA plates (Costar), washed with Tris-buffered Saline+0.05%Tween, blocked using 2% w/v Bovine serum albumin (BSA; Sigma) and incubated with pleural lavage fluid overnight at 4°C prior to detection of bound IgM with a goat anti-mouse IgM-HRP (1020-05 Southern Biotech) for 1h at 37°C. 50μl of 3,3 ‘,5,5 ‘-Tetramethylbenzidine (TMB; Biolegend) was added, allowed to develop and stopped with 50μl 0.18M H_2_SO_4_. Plates were read at 450nm on Tecan Infinite 200PRO plate reader.

### Statistical analysis

Power calculations showed that for our most commonly measured parameters (cell number) 6 mice per group provide sufficient power (90%) to detect at least a 1.5-fold difference between the groups, which we regard as an acceptable cut-off for identifying important biological effects. No randomization and no blinding was used for the animal experiments. Whenever possible, the investigator was partially blinded for assessing the outcome (confocal analysis). All data were analysed using Prism 9 (Graphpad Prism, La Jolla, CA, USA). Statistical tests performed, sample size and number of repetitions for each data set, are described within the relevant figure legend.

## Results & discussion

FALCs are key sites of immune orchestration within the serous cavities, in order to determine the early immune response to pristane within the cavities we first quantified key immune cell populations within FALC containing adipose depots (omentum, pericardium and mediastinum) of both the peritoneal (site of delivery) and pleural (distal site) cavities. As we were interested to better understand the initiation of inflammation within the pleural cavity, we utilised C57BL/6 mice that have been shown to develop pulmonary complications following delivery of pristane. To determine the role of FALCs in co-ordinating the immune response within the body cavities following pristane exposure we injected a low volume (300ul) of pristane into the peritoneal cavity (to limit potential for direct leakage of pristane from the peritoneal into the pleural cavity) and isolated the omental and pericardial adipose tissues after 3 days.

Weighing of the tissues revealed an increase in omental but not pericardial adipose weight (Figure 1A), this increase in weight appeared to be mediated by an expansion in the size of FALCs within the tissue as determined via analysis of cluster perimeter using Image J following whole-mount immuno-fluorescence analysis of fixed tissues (Figure 1B). To determine which immune cells may be responsible for the early differential expansion of FALCs within the omentum and pericardium following pristane exposure we analysed the % area of expression with individual FALCs via whole-mount immuno-fluorescence staining for IgM (B-lymphocytes, red), F4/80 (serous cavity macrophages, yellow) and Ki67 (proliferation, magenta) (Figure 1C). There was a significant reduction in the % area of IgM within clusters of the omentum at day 3 following pristane delivery, suggesting either a reduction in the number of B cells present within the tissue or an indication that the B cells within this tissue had undergone class switch to express a different antibody isotype (Figure 1D). In contrast, there was no significant difference in the percentage area of IgM expression within the pericardial FALCs after pristane delivery (Figure 1C-D). This data suggested that B cells were not the main cell type responsible for early FALC expansion within the omentum. Peritoneal macrophages have been shown to migrate to the omentum during infection of the peritoneum (Christian et al. 2022; Zhang et al. 2019; Jones et al. 2015) as such, we also assessed F4/80 within FALCs of the omentum and pericardium and found no significant increase in the percentage area of expression, of this serous cavity macrophage receptor at 3 days after pristane injection (Figure 1D). Percentage area of expression of F4/80 was variable mouse to mouse and there was a trend in some experiments towards an increase in macrophage recruitment within the omentum but this trend was not seen in the pericardium. In contrast to these cell-type defining markers, we saw a significant increase in percentage area of expression of both omental and pericardial FALCs for the nuclear cell cycle protein Ki67 which is an accepted readout of cellular proliferation (Figure 1 C-D).

**Figure 1.**
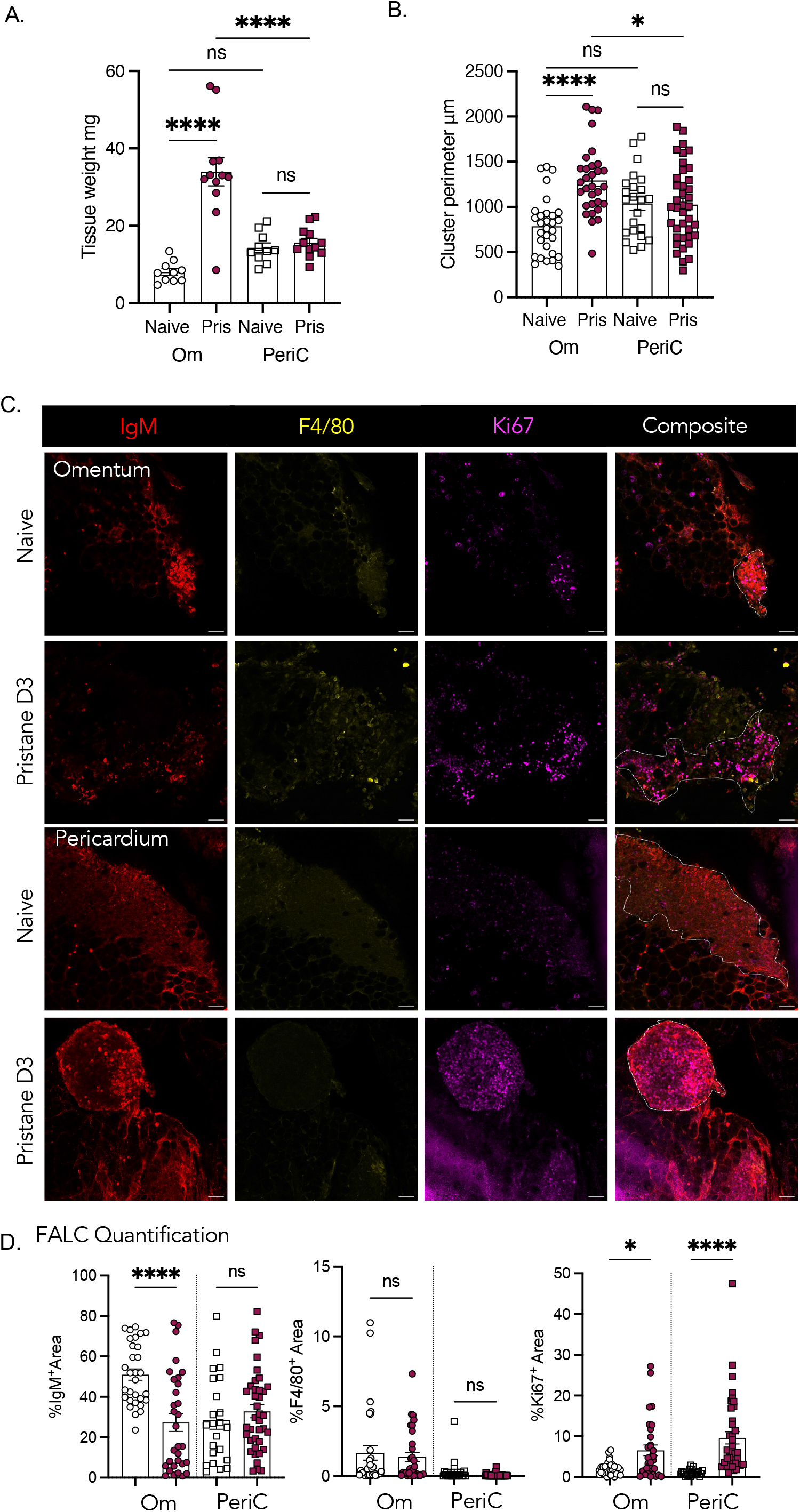
FALCs are activated in response to intra-peritoneal pristane delivery. C57BL/6 mice were injected i.p. with 300μl pristane or left naive, 3 days later omentum & pericardium were isolated and weighed (A), whole-mount immuno-fluorescence confocal microscopy was undertaken on omenta and pericardia to determine cluster perimeter (B), IgM (Red), F4/80 (Yellow) and Ki67 (magenta) expression (C), area of expression per cluster was quantified using Image J (D). Data in A pooled from three independent experiments, n=3-5 mice per group. Data in B & D pooled from two independent experiments, data points represent individual FALCs from 3-5 mice per group. Data in C representative of 2 experiments, n=3-5 mice per group. Student’s T-Test, *=P <0.05, **** P=<0.0001, ns = non-significant, scale bar = 50μM.

To further confirm which cells are responsible for expanded cluster size within the omentum following pristane delivery we assessed using multi-parameter flow cytometry the presence of CD45^+^CD19^+^MHCII^+^CD11b^+^ B1 B cells, CD45^+^CD19^+^MHCII^+^CD11b^-^B2 B cells, CD45^+^CD19^-^Ly6G^-^SigF^+^SSC-A^hi^ Eosinophils, CD45^+^CD11b^+^Ly6G^+^ neutrophils, CD45^+^CD19^-^Ly6G^-^SigF^-^TCRβ^-^Ly6C^+^ monocytes and CD45^+^CD19^-^Ly6G^-^TCRβ^-^SigF^-^Ly6C^-^CD11b^+^F4/80^+^Tim4^+/-^macrophages within peritoneal exudate cells (PEC), digested omentum, pleural exudate cells (PLEC) and digested pericardium of naïve mice and those exposed 3 days earlier to 300μl of intra-peritoneal pristane. There was a trend towards reduced numbers of both B1 & B2 B cells within the PEC with a significant increase found within the omentum (Figure 2A), in contrast to the peritoneal cavity, there was a significant increase in the numbers of B1 and B2 B cells within pleural lavage at day 3 following pristane injection but no corresponding increase in the pericardium. This data suggests that B cells contribute to differential expansion of FALCs following pristane delivery. The increase in B cells within the omentum was not sufficient to account for the increased cluster size noted during perimeter analysis (Figure 1B) however we noted a striking increase in the number of eosinophils and neutrophils within the omentum but not the pericardium (Figure 2B). We have previously shown that the omentum is a key site for neutrophil recruitment during peritonitis (Jackson-Jones 2020), to investigate whether the mechanism via which neutrophils are recruited to omental FALCs after pristane exposure may be similar to that during peritonitis we assessed the release of the neutrophil recruitment chemokine CXCL1 and found significantly increased CXCL1 release from the omentum during 2h of culture when isolated from pristane exposed compared to naïve mice (Figure 2C). In contrast, there was no increased CXCL1 release from the pericardium and no significant recruitment of neutrophils to this site suggesting that CXCL1 release may be a factor in the recruitment of neutrophils to the omentum in this model. There was a small increase in the number of monocytes within the PEC at day 3 following pristane delivery and a trend towards an increase in the omentum but no measurable difference in number within the PLEC or pericardium (Figure 2D). The macrophage disappearance reaction is a phenomenon that characterises the stark absence of large cavity macrophages within the serous cavities rapidly following induction of inflammation (Barth et al. 1995). Using a combination of CD11b, F4/80 and Tim4 we were able to define two major populations of macrophages that could be detected in all four sites of interest. As reported previously, the majority of F4/80 high large cavity macrophages (LCM) within naive serous cavities were found to be Tim4^+^, this population was absent in the peritoneal cavity at day 3 after pristane delivery and replaced by an expanded Tim4^-^ population (Figure 2E-F). A small increase in Tim4^+^ macrophages was found within the omentum but this did not reach significance whereas a large increase in Tim4^-^ macrophages was seen at this site (Figure 2F) mirroring previous studies that found Tim4^-^ cells at this site be of bone marrow origin (Etzerodt et al. 2020), as well as omental accrual of equivalent CD102^-^ inflammation-elicited macrophages (Louwe et al. 2022). A small but significant increase in the number of Tim4^+^ LCM within the pleural cavity was found at 3 days after pristane delivery (Figure 2D). It does not appear that the increase in the number of Tim4^+^ macrophages found within the pleural cavity is due to their local proliferation as analysis of the cell cycle protein Ki67 within Live, CD45^+^CD19^-^CD11b^+^ cells that are MHCII low shows no significant increase in expression at either the population or per cell level (data not shown). It would be interesting to investigate in future experiments whether any of the peritoneal macrophages that are lost following pristane injection migrate via the diaphragm to account for this increase found in the pleural fluid. Loss of some of the Tim4^+^ cells within the peritoneal space following pristane injection can be accounted for via migration to the omentum however this by no means accounts for the total loss, indeed as shown in *E*.*coli* infection experiments (Zhang et al. 2019) we also saw clots of cells or granulomas within the cavity at this time point, however we have not yet investigated the cellular composition of such.

**Figure 2.**
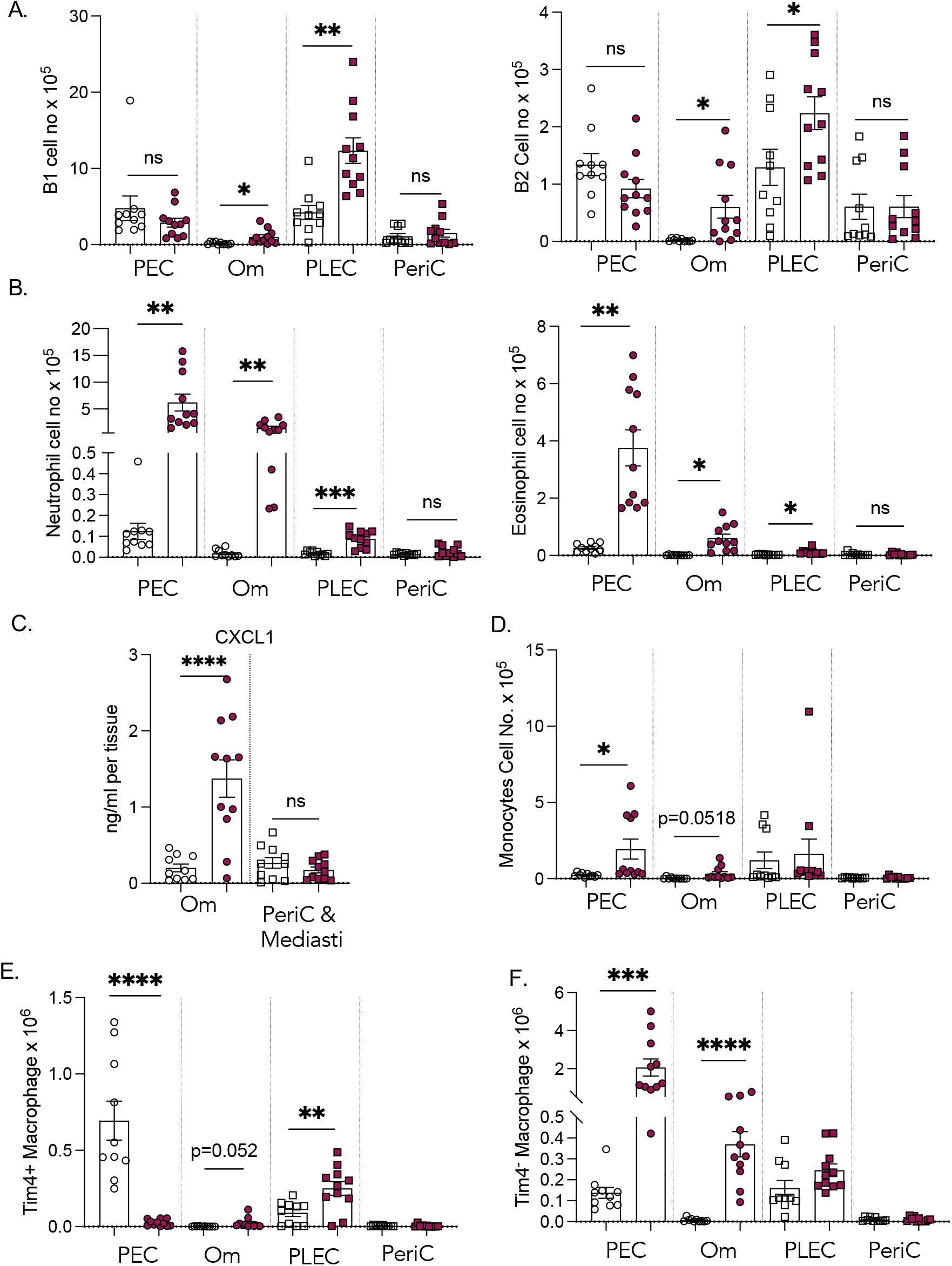
Pristane delivery induces differential immune cell changes within the peritoneal and pleural cavities. C57BL/6 mice were injected i.p. with 300μl pristane (maroon) or left naïve (white), 3 days later Peritoneal lavage, Pleural lavage, omentum, pericardium and mediastinum were isolated and tissues weighed. Omenta and pericardium were digested and assessed by flow cytometry alongside PEC & PLEC (A-B, D-E) for the presence of immune cells (Live, singlets, CD45^+^) including B1 B cells (Ly6G^-^CD19^+^MHCII^+^CD11b^+^), B2 B cells (Ly6G^-^CD19^+^MHCII^+^CD11b^-^), Neutrophils (Ly6G^-^), Eosinophils (Ly6G^-^SigF^+^SSCA^hi^), Monocytes (Ly6G^-^CD19^-^TCRβ^-^SigF^-^Ly6C^+^), Tim4+ Macrophages (Ly6G^-^CD19^-^ TCRβ^-^SigF^-^Ly6C^-^F4/80^+^CD11b+Tim4^+^),Tim4-Macrophages (Ly6G^-^CD19^-^TCRβ^-^SigF^-^Ly6C^-^F4/80^+^CD11b+Tim4^-^). Omenta and mediastina were placed in culture at 37°C 5 %CO_2_ for 2h and total amount of CXCL1 secreted into the culture supernatant within 2h was quantified using a legendplex array, total CXCL1 secreted from whole FALC containing tissue was then calculated using whole tissue weights within each cavity (C). Data pooled from 3 independent experiments n=3-5 mice per group per experiment, Student’s T-Test, *=P <0.05,**=P<0.01,^***^ P=<0.001,****P=<0.0001., Student’s T-Test, *=P <0.05, **=P <0.01,***P=<0.001,**** P=<0.0001.

We next assessed the phenotype of B1 B cells within the peritoneal and pleural cavities and their integral FALC containing adipose tissues. B1 cells from the PEC, digested omentum, PLEC and digested pericardium of naïve mice and those exposed 3 days earlier to 300μl of intra-peritoneal pristane were assessed via flow cytometry to determine nuclear levels of Ki67 (Figure 3A) and the plasma cell surface marker CD138 (syndecan-1) (Figure 3B). There was a significant increase in the expression of Ki67 by B1 cells within the PEC, PLEC and pericardium at 3 days after pristane exposure, indicating that these cells were actively progressing through cell cycle. B1 cell proliferation may account for the increased Ki67 detected within the pericardium following pristane exposure (Figure 1C & D). B1 cells within PLEC and pericardium were also found to have increased expression of CD138 at 3 days following pristane exposure, suggesting that these cells may be actively secreting antibodies (Figure 3B). This finding was in contrast to B1 cells of the PEC which had reduced expression of CD138 and the omentum in which no difference in CD138 expression was found following pristane exposure (Figure 3B). We next placed omentum and mediastinum in culture and quantified the secretion of antibodies from the tissues per mg in 2h to enable calculation of the total amount of antibody released from the total pleural adipose tissues. There was significantly increased secretion of IgM, IgG2a, IgG1 and a trend for increased IgA from pleural FALC containing tissues at day 3 following pristane exposure (Figure 3C). IgA was only detected in one of the naïve culture supernatants making statistical analysis unviable. Increased secretion of total IgM, IgG2a and IgG1 was in contrast to the omentum which did not significantly increase total secretion of these antibody isotypes. Pristane has been shown to cause lymphocyte apoptosis both *in vitro* and *in vivo* (Calvani et al. 2005), as such reduced antibody secretion by the omentum (the FALC containing adipose at the site of injection) may in part be due to the loss of viable B cells within this tissue, as supported by reduced IgM staining (Figure 1D), and no increase in CD138 by flow cytometry. Furthermore, the striking difference in neutrophil numbers within the omentum after pristane delivery in contrast to the failed recruitment to the pericardium & mediastinum (Figure 2B) may modify the functional capacity of the tissue to secrete antibodies within the 2h window assessed.

**Figure 3.**
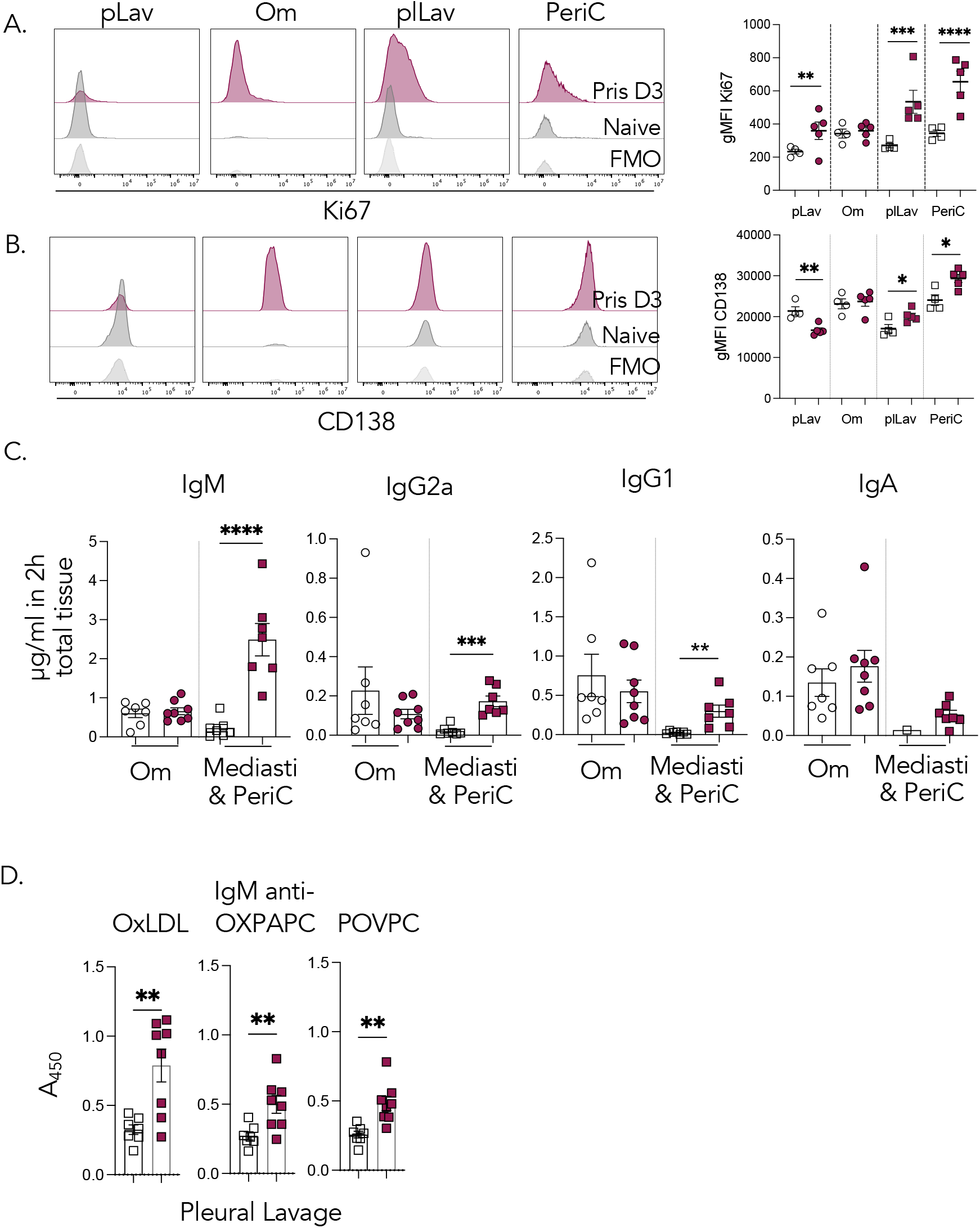
Pleural FALCs produce antibodies in response to intra-peritoneal pristane delivery. C57BL/6 mice were injected i.p. with 300μl pristane or left naive, 3 days later Peritoneal lavage, Pleural lavage, omentum, pericardium and mediastinum were isolated and tissues weighed. Omenta were dissected into two and each half was weighed; half of the omentum and pericardium were digested and assessed by flow cytometry alongside PEC & PLEC (A&B) for the expression of Ki67 and CD138 within B1 B cells (Live, singlets, CD45+Ly6G^-^CD19^+^MHCII^+^CD11b^+^). Omenta and mediastina were placed in culture at 37°C 5 %CO_2_ for 2h and total amount of IgM, IgG2a, IgG1 and IgA secreted into the culture supernatant within 2h were quantified using a legendplex array, total antibody amount secreted from whole FALC containing tissue was then calculated using whole tissue weights within each cavity (C). Cell free pleural lavage was assessed for presence of anti-phospholipid IgM antibodies via ELISA (D). Data pooled from two independent experiments n=3-4 per group, only one naïve mediastinum culture sample had detectable IgA. Student’s T-Test, *=P <0.05, **=P <0.01, *** P=<0.001, **** P=<0.0001.

Phospholipid antibodies are implicated in the pleural pathogenesis of SLE, as such we next assessed whether we could detect anti-phospholipid antibodies within the pleural fluid at day 3 after exposure to pristane. As IgM was the antibody most significantly increased at this timepoint in pleural adipose culture supernatants and IgM having been implicated in the pathology of DAH in C57BL/6 mice we determined via ELISA the presence of IgM recognising the naturally occurring phospholipids oxidised low-density lipoprotein (oxLDL), oxidised-1-palmitoyl-2-arachidonoyl-*sn*-glycero-3-phosphorylcholine (oxPAPC) and 1-palmitoyl-2-(5 ‘-oxo-valeroyl)-*sn*-glycero-3-phosphocholine (POVPC) within pleural lavage fluid. IgM recognising all three phospholipid species could be detected at day 3 following intra-peritoneal pristane exposure (Figure 3D) highlighting the local presence of anti-phospholipid antibodies within the pleural cavity early during initiation of lupus pleuritis in a murine model.

Collectively, our findings highlight a role of pleural FALCs of the pericardium and mediastinum and the local production of IgM auto-antibodies in the aetiology of lupus-like pleuritis in a C57BL/6 mouse model.

## Author contributions

KB, LG, AH, BE, Investigation, writing-review. SM, NI, Investigation, supervision, writing-review. **LHJJ** Investigation, Formal Analysis, Funding Acquisition, Supervision, Conceptualization, Project administration, Writing-original draft, visualization.

## Acknowledgements

this work was funded by a Wellcome Seed award in science (213697/Z/18/Z) and an MRC NIRG (MR/V031767/1) both to LJJ. We would like to thank members of the microbes, pathogens and immunity theme at Lancaster University for useful discussions about the work.

## Figure Legends

**Supplementary table 1.** Antibodies used in the study.

| Antigen Name | Conjugate | Clone | Manufacturer |
| --- | --- | --- | --- |
| IgM | HRP |  | Southern Biotech |
| CD11b | PE-Dazzle CF594 | M1/70 | Biolegend |
| CD19 | BV421 | 6D5 | Biolegend |
| Siglec F | BV421 | E50-2440 | BD |
| TCRb | BV421 | H57-597 | Biolegend |
| CD45 | BV650 | 104 | Biolegend |
| F4/80 | PE/Cy7 | BM8 | Biolegend |
|  | PE/Cy7 | BM8 | Invitrogen |
| Ki-67 | FITC | REA183 | Miltenyi Biotech |
| Ly6C | Alexa-Fluor 700 | HK1.4 | Biolegend |
| Ly6G | FITC | REA526 | Miltenyi Biotech |
| MHCII IIA/IE | APC/Fire™ 750 | M5/114.15.2 | Biolegend |
| Tim4 | PE | RMT4-54 | Biolegend |
| F4/80 | FITC | BM8 | Biolegend |
| Donkey anti-mouse IgM | Rhodamine Red X | Polyclonal | Jackson Laboratories |
| Ki-67 | VioRed 667 | REA183 | Miltenyi Biotech |
| Supplementary table 1. Antibodies used in the study |  |  |  |

**Supplementary Figure 1.**
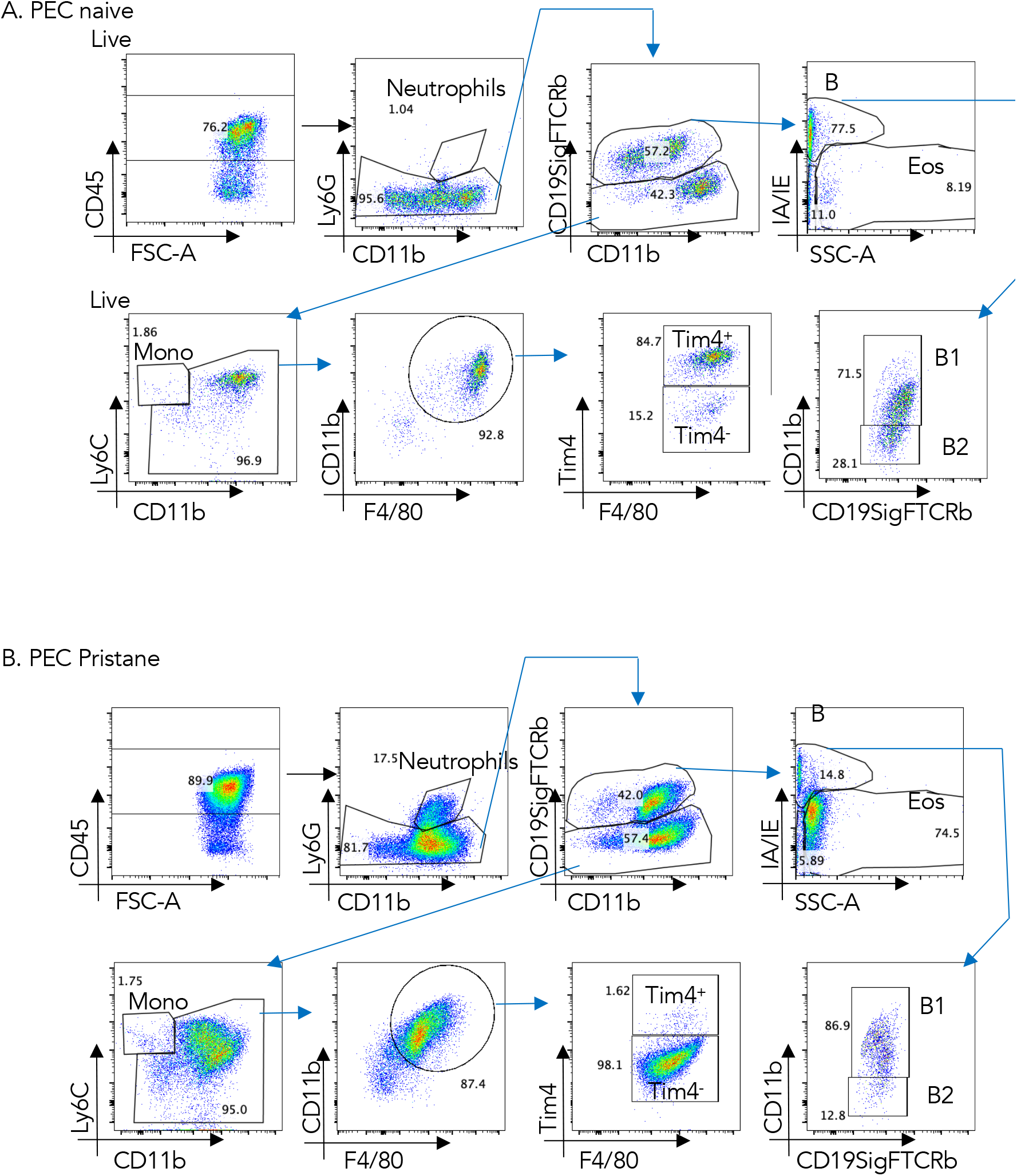
Representative flow cytometry gating strategy.

